# Behavioural and transcriptomic characterization of the comorbidity between Alzheimer’s disease and Major Depression

**DOI:** 10.1101/2020.07.31.231159

**Authors:** Ana Martín-Sánchez, Janet Piñero, Lara Nonell, Magdalena Arnal, Elena M. Ribe, Alejo Nevado-Holgado, Simon Lovestone, Ferran Sanz, Laura I. Furlong, Olga Valverde

**Affiliations:** Neurobiology of Behaviour Research Group (GReNeC-NeuroBio), Department of Experimental and Health Sciences, Universitat Pompeu Fabra; Neuroscience Research Program, IMIM-Hospital del Mar Research Institute, Barcelona, Spain; Research Programme on Biomedical Informatics (GRIB), IMIM (Hospital del Mar Medical Research Institute), Universitat Pompeu Fabra, Barcelona, Spain; MARGenomics core facility, IMIM (Hospital del Mar Medical Research Institute), Barcelona, Spain; Department of Psychiatry, University of Oxford, Oxford OX3 7JX, UK; Oxford Health NHS Foundation Trust, Oxford OX3 7JX, UK; Janssen-Cilag, UK

## Abstract

Major Depression (MD) is the most prevalent psychiatric disease in the population and is considered a prodromal stage of the Alzheimer’s disease (AD). Despite both diseases having a robust genetic component, the common transcriptomic signature remains unknown. In this regard, we investigated the cognitive and emotional responses in 3- and 6-month-old in APP/PSEN1-Tg mutant mice, before β-amyloid plaques were detected. Then, we studied the deregulation of genes and pathways in prefrontal cortex, striatum, hippocampus and amygdala, using transcriptomic and functional data analysis. The results demonstrated that depressive-like and anxiety-like behaviours, as well as memory impairments are already present at 3-month-old together with the deregulation of several genes and gene sets, including components of the circadian rhythms, electronic transport chain and neurotransmission. Finally, DisGeNET GSEA provides translational support for common depregulated gene sets related to MD and AD. Altogether, the results demonstrate that MD could be an early manifestation of AD.

## INTRODUCTION

Depressive disorder affects over 4.4% of the population^1^, and it is characterized by feelings of sadness, anhedonic state, changes in appetite and sleep, feelings of worthlessness, guilt, and may lead to attempts of suicide^1^. Major Depression (MD) manifests throughout the life course. In older people it is often associated with poor cognition that may not return to normality with effective treatment of the mood disorder^2,3^. In addition to the cognitive phenotype of MD in older people, a substantial body of data strongly suggests onset of MD in later life is associated with increased risk of Alzheimer’s disease (AD), the commonest form of dementia^4–6^. Furthermore, although cognitive and functional impairments are the predominant symptoms of AD, other behavioural and psychological symptoms of dementia (BPSD) that include depression, sleep and activity disturbance are common manifestations of the disease^7,8^. These observations have led to the suggestion that MD may in some cases be a prodromal phase of AD. Indeed, both MD and AD have a considerable heritable component and share some mechanistic components including neuroinflammation^9,10^, oxidative stress^11^, certain dysregulations in cellular signalling^12^ and neurotransmission^13,14^ amongst others.

Despite these advances, the aetiology of the comorbidity between AD and MD remains unknown although the observation that multiple behaviours that are reminiscent of BPSD are also observed in rodent models of disease^15^ suggesting that there might have common molecular processes. Hereby, we study for the first time, using behavioural, transcriptomic and bioinformatic approaches, depressive-like symptoms in an AD mouse model (APP/PSEN1-Tg mice) at ages before the neuropathological features are manifest and when cognitive impairment is not evident^16^. To that end, we have performed RNA sequencing analyses to study changes that occur in brain areas related to the control of behavioural and cognitive behaviours, including prefrontal cortex (PFC), striatum, hippocampus and amygdala, in mice at 3 and 6 months old. In order to study the mechanisms involved in the AD-MD comorbidity, a pre-ranked Gene Set Enrichment Analysis (GSEA) has been carried out. Using the transcriptomic results, we evaluated the enrichment in genes sets obtained from public resources containing functional and disease information.

## RESULTS

The transgenic mice expressing human APP/PSEN1-Tg carrying familial AD mutations used in these studies (B6C3-Tg) have an onset of plaque pathology at around 6 months, which is followed by cognitive impairments with both increasing by 12-15 months of age. To explore behavioural alterations in these mice, we used a range of well-established experimental paradigms at 3 months (before the onset of pathology) and 6 months (at onset). Then, we evaluated the differentially expressed genes and pathways in PFC, striatum, hippocampus and amygdala at both ages.

### Anxiety-related behaviour in APP/PSEN1-Tg mice

To evaluate if APP/PSEN1-Tg animals experience anxiety-like behaviour, 3 and 6-month-old APP/PSEN1-Tg were subjected to the elevated plus maze (EPM). The APP/PSEN1-Tg transgenic mice spent lower percentage of time in open arms than Non-Transgenic at both 3 (t_30_= 2.30, p=0.0286; Figure 1A) and 6 months old (t_26_=2.419, p=0.0229; Figure 1D), whereas no differences were found in the total number of arm entries (t_30_= 0.65, p=0.520, Figure 1B and t_26_=1.21, p=0.236, Figure 1E), indicating that APP/PSEN1-Tg animals displayed higher level of anxiety-like behaviours without locomotor impairments.

**Figure 1.**
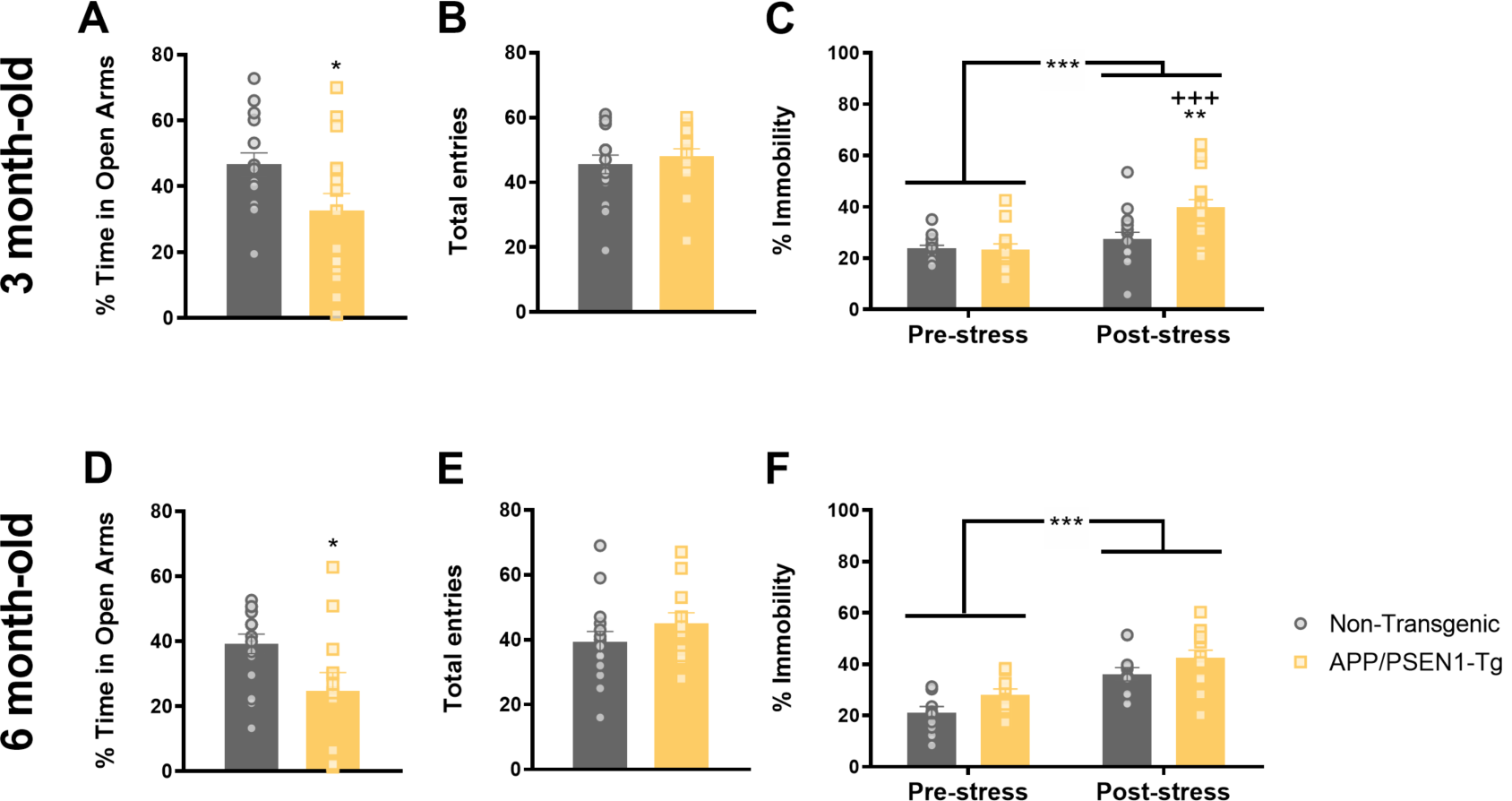
APP/PSEN1-Tg animals show early anxiety-like and despair-like behaviours induced by stress. Panels A,B,D and E represent the EPM results, and panels C and F correspond to the TST results. Grey (Non-Transgenic) and yellow (APP/PSEN1-Tg) bars represent the percentage of time spent in open arms (A, D) and total number of entries (B,E) at 3 (n=16 per group) and 6 months (n=14 per group). (*p < 0.05, Student’s t-test Non-Transgenic *vs* APP/PSEN1-Tg). Grey (Non-Transgenic) and yellow (APP/PSEN1-Tg) bars represent the percentage of time that animals are immobile in TST at 3 (n=16 per group) (C) and 6 months-old (Non-Transgenic n=12, APP/PSEN1-Tg n=14) (F) in pre-stress and post-stress conditions. Data are presented as mean ± SEM. ***p<0.001 Pre-stress *vs* post-stress condition; +++ p<0.001 comparison APP/PSEN1-Tg pre-stress *vs* post-stress, **p<0.01 comparison APP/PSEN1-Tg *vs* Non-Transgenic post-stress condition. (two-way ANOVA).

### Early onset of despair-like behaviour in APP/PSEN1-Tg mutant mice

Stress induced immobility is a despair responsive phenotype often used as a proxy for mood state in rodent models. To evaluate this response, we performed the tail suspension (TST) in APP/PSEN1-Tg transgenic and control mice.

At 3 months old, the analysis of the TST results using an ANOVA for repeated measurements revealed *stress* (F_1,30_ =20.77, p<0.001), *genotype* (F_1,30_=5.27, p=0.029) and *stress×genotype* (F_1,30_=8.61, p=0.006; Figure 1 C) effects. After Bonferroni’s correction, results showed that transgenic mice presented higher stress-induced immobility (p<0.001) in comparison with the pre-stress condition. Stress only increased the percentage of the immobility in APP/PSEN1-Tg mutants in comparison with Non-Transgenic animals (p=0.005).

At 6 months, the ANOVA for repeated measurements showed *stress* (F_1,24_ =31.33, p<0.001) and *genotype* (F_1,24_=4.62, p=0.042; Figure 1F) effects. These results indicate that transgenic mice spent a higher percentage of immobility time and after repeated stress, but also this behaviour increases in both groups of mice.

### Early-adulthood long-term memory impairments in APP/PSEN1-Tg mice

Loss of memory is affected early during the time-course of the AD, being one of the most recognizable symptoms of the disease^17^. For this reason, we assessed the short- and long-term recognition memory in 3- and 6-month-old APP/PSEN1-Tg mice using the novel object recognition test.

At 3 months old, a two-way ANOVA showed a *genotype* (F_1,29_=8.30, p=0.007), and *object* effect (F_1,29_=36.93, p=0.001; Figure 2A). The *genotype* effect indicates that control animals had greater discrimination index than APP/PSEN1-Tg group.

**Figure 2.**
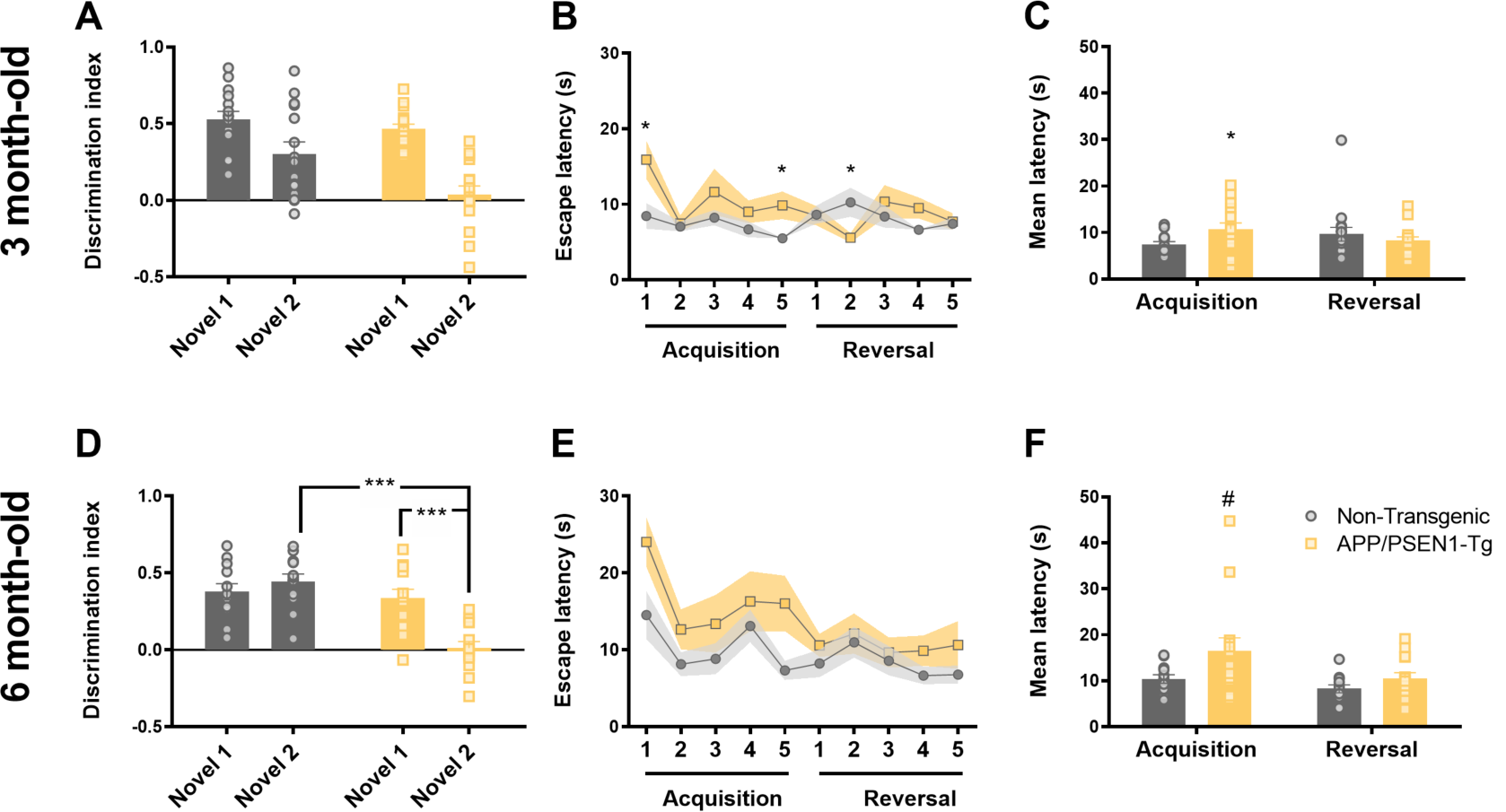
The onset of memory impairments in APP/PSEN1-Tg occurs at 3months-old. Novel object recognition test: Grey (Non-Transgenic) and yellow (APP/PSEN1-Tg) bars represent the novel object 1 and 2 discrimination index (%) at 3 (A) (Non-Transgenic n=15, APP/PSEN1-Tg n=16) and 6 months old (D) (Non-Transgenic n=12, APP/PSEN1-Tg n=14). Data are presented as mean ± SEM. *p<0.05, **p < 0.01 (two-way ANOVA). T-maze left-right discrimination learning: (B, E) Solid (Non-Transgenic) and dotted line (APP/PSEN1-Tg) represent the escape latencies of both groups at two different ages. (ANOVA of repeated measurements *p<0.05; +p<0.05 comparison APP/PSEN1-Tg *vs* the 1^st^ trial; *p<0.05 comparison APP/PSEN1-Tg *vs* Non-Transgenic). (C, F) Grey (Non-Transgenic) and yellow (APP/PSEN1-Tg) bars represent the mean ± SEM trials to criterion during acquisition and reversal phases latencies at 3 (n=16 per group) and 6 months-old (Non-Transgenic, n=12, APP/PSEN1-Tg n=14). (F) At 6 months-old, transgenic mice tend to reach the criterion later than control groups during acquisition. (Student’s t-test, Non-Transgenic *vs* APP/PSEN1-Tg, *p<0.05; #p=0.068).

At 6 months, a two-way ANOVA revealed a main effect of *genotype* (F_1,24_=22.44, p<0.001), an *object* effect (F_1,24_=6.51, p=0.018) and *object × genotype* interaction (F_1,24_=14.64, p=0.001; Figure 2D). The *genotype* effect shows that control animals had greater discrimination index than transgenic mice. The *post-hoc* Bonferroni’s analysis showed that Non-Transgenic mice could discriminate both the novel object 1 and 2 in the same manner (p=0.394), whereas APP/PSEN1-Tg animals could only discriminate novel object 1, having a greater discrimination index for the novel object 1 than novel object 2 (p<0.001). Non-Transgenic animals had higher discrimination index than APP/PSEN1 when they are exposed to novel object 2 (p<0.001). In fact, APP/PSEN1-Tg animals could not discriminate between familial and novel object 2, indicating important memory impairments.

### Early poor left-right discrimination learning of APP/PSEN1-Tg mice

In order to evaluate working memory and attention deficits present in AD^18^, we performed a left-right discrimination learning paradigm in 3 and 6 month APP/PSEN1-Tg transgenic and Non-Transgenic mice using the T-maze.

The two-way ANOVA showed a *trial* effect (F_9,270_= 2.152, p=0.026) and *trial × genotype* (F_9,270_=2.023 p=0.037; Figure 2B) at 3 months old. The *post-hoc* analysis revealed that APP/PSEN1-Tg animals took more time to reach de platform than control animals on trial 1 (p=0.044) and trial 5 (p=0.027) during the acquisition learning. However, this analysis suggests that control mice spent more time to reach the platform during the trial 2 of the reversal-learning phase (p=0.02) than transgenic mice. The pair-wise comparison showed that APP/PSEN1-Tg reduced the time to escape on acquisition trial 2 (p=0.037) and trial 2 of the reversal learning (p=0.017) in comparison with the trial 1 of the acquisition. We found that APP/PSEN1-Tg animals spent more trials than Non-Transgenic animals (t_30_=2.35, p=0.026; Figure 2C) to reach the acquisition criterion. Nevertheless, both groups spent a similar number of trials to reach the criteria during reversal learning (p=0.443).

At 6 months old, the ANOVA revealed a *trial* (F_9,216_= 3.84, p<0.001) and *genotype* (F_1,24_=6.53, p=0.017, Figure 2E) effects, suggesting that APP/PSEN1-Tg mutant mice showed longer latencies to reach the platform than control animals during acquisition and reversal learning phases. The mean latency to get the acquisition criteria showed a trend to be statistically different in the number of trials that APP/PSEN1-Tg in comparison with control mice (t_24_=1.91; p=0.068; Figure 2F).

### Hyposmia in APP/PSEN1-Tg mice at 6 months old

Since olfactory impairment is present in up to 90% of AD patients^19^, we assessed the olfactory function in transgenic and control mice (Figure 3A and C).

**Figure 3.**
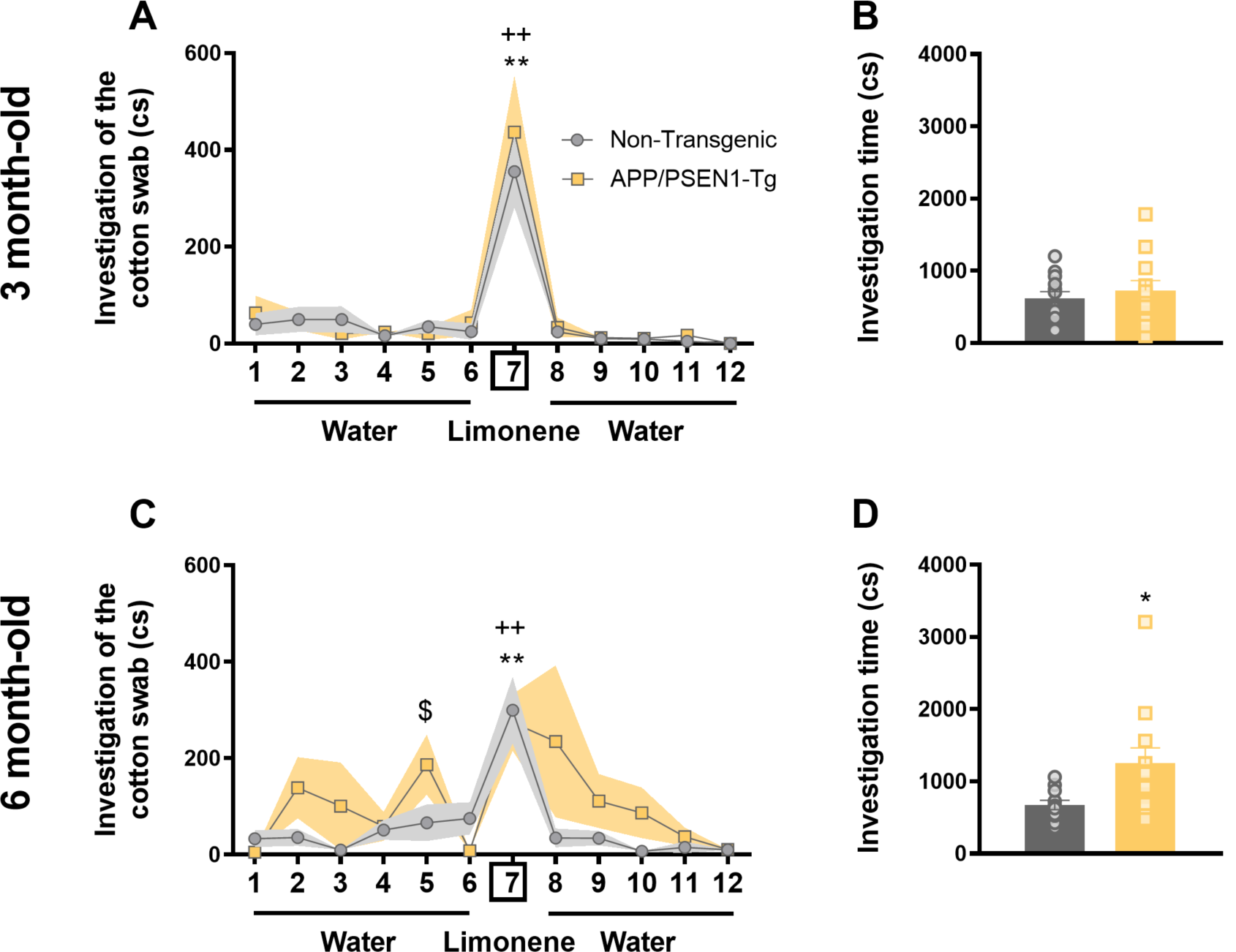
APP/PSEN1-Tg mice show olfaction disruptions at 6 months old. Graphs show the time (centiseconds, cs) that APP/PSEN1-Tg and Non-Transgenic animals spent investigating a cotton swab through consecutive one-minute presentations (mean± SEM) of distilled water (presentations 1-6 and 8-12) and limonene (presentation 7) at 3 (A) and 6 months (C). The exploration time during the first presentation of the cotton swab with water [1] was compared with that of the first presentation of limonene [7] in each group. (Wilcoxon test, **p<0.01 comparison APP/PSEN1-Tg water *vs* limonene presentations; ++p<0.01 comparison Non-Transgenic water *vs* limonene presentations; Kruskal-Wallis test $p<0.05 comparison APP/PSEN1-Tg *vs* Non-Transgenic). Grey (Non-Transgenic) and yellow (APP/PSEN1-Tg) bars represent the total investigation time at (B) 3 (n=12 per group) and 6 (D) months old (Non-Transgenic n=11, APP/PSEN1-Tg n=12) (Student’s t-test *p<0.05). Data are presented as mean ± SEM.

At 3 months old, both experimental groups could discriminate between the first water presentation and limonene odour (Wilcoxon test, Non-Transgenic group, p=0.05 and APP/PSEN1-Tg, p=0.01; Figure 3A).

At 6 months, both APP/PSEN1-Tg transgenic mice and Non-Transgenic mice could discriminate between the first water presentation and limonene (Wilcoxon test, Non-Transgenic, p=0.006, and APP/PSEN1-Tg, p=0.008; Figure 3C). Additionally, APP/PSEN1-Tg mutant mice showed a higher investigation time during the fifth water presentation than control group (Kruskal-Wallis test, χ^2^=3.84, p=0.05; Figure 3C). Whereas both experimental groups spent the same time investigating the cotton swab at 3 months old (t_22_=0.66, p=0.52; Figure 3B), APP/PSEN1-Tg mutant increased the investigation time than Non-Transgenic control animals, showing an indiscriminate investigation at 6 months-old (t_21_=2.52, p=0.02; Figure 3D), independently of the type of odour presented.

### DE analysis in PFC, striatum, hippocampus and amygdala at two different ages

We performed gene expression analysis in regions of the brain known to be loci for these functions. We compared the transcriptome signature of APP/PSEN1-Tg transgenic and control mice at 3 and 6 months old, to determine the significant differentially expressed (DE) genes with a cut-off |logFC | < 0.585, adjusted p<0.05, deregulated in the brain areas of interest (Figure 4; Supplementary Tables 2-5).

**Figure 4.**
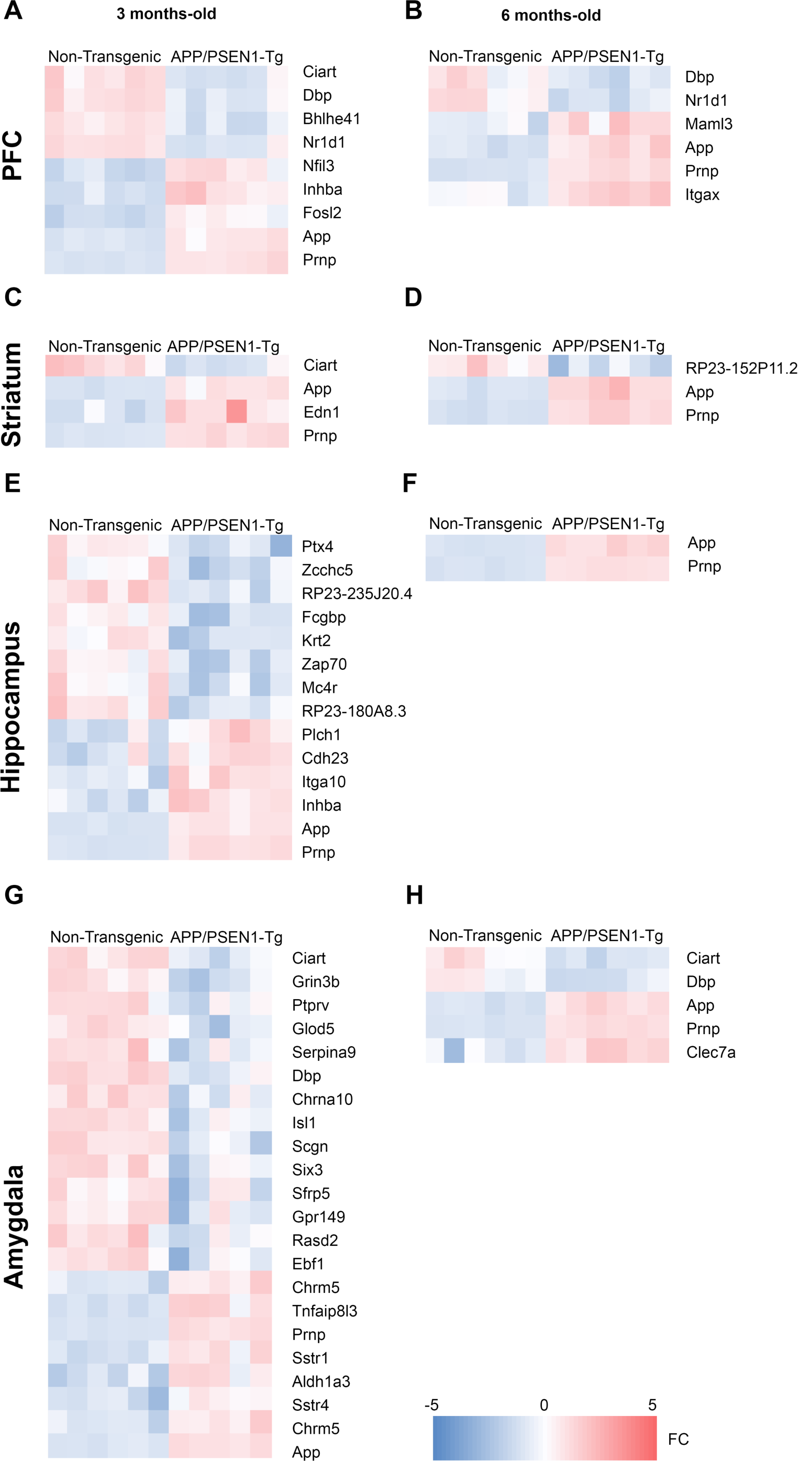
Differentially expressed genes in the PFC, striatum, hippocampus and amygdala from APP/PSEN1-Tg mice at 3 and 6 months-old. Heatmaps representing the degree of change for the differentially expressed genes at 3 (A,C,E,G) and 6 (B,D,F,H) months between control and APP/PSEN1-Tg (5-6 independent brain samples per area). (G) The heatmap from amygdala at 3 month-old is a reduced representation from >100 up and downregulated genes (p < 0.05;|logFC|>0.59). Legend (bottom) indicates the color-coded fold-change scale (−5<FC<5) where negative values represen downregulation in blue, and positive upregulation in red.

Among the dysregulated genes in PFC of younger APP/PSEN1-Tg mice, we found 5 corresponding to the so-called canonical clock genes: *Ciart, Dbp, Bhleh41* and *Nr1d1* were downregulated, and *Nfil3* was upregulated, together with other genes, such as *App, Prpn, Inhba*, and *Fosl2* (Figure 4A; Supplementary Table 2). At 6 months-old mice, *Dbp* and *Nr1d1* remained downregulated in this area, whereas *App, Prnp* were also upregulated together with *Itgax* and *Maml3* (Figure 4B; Supplementary Table 2).

In the striatum of 3-months-old mice, we only found the downregulation of *Ciart*, and the overexpression of *App, Prpn* and *Edn1* (Figure 4C; Supplementary Table 3). By contrast, the deregulation of older mice in the striatum was restricted to only one downregulated transcript, *RP23-152P11*.*2*, whilst *App* and *Prpn* remained upregulated (Figure 4D; Supplementary Table 3).

In the hippocampus of 3-months-old APP/PSEN1-Tg mice, 8 genes were downregulated, including *Ptx4* and *Mc4r* and *Zap70*. Among the upregulated genes, we found *Plch1, Cdh23, Itga10, App, Prnp*, and *Inhba* (Figure 4E; Supplementary Table 4). At 6 months, the analysis revealed that only *App* and *Prnp* remained upregulated (Figure 4F; Supplementary Table 4).

Interestingly, the amygdala of young transgenic mice was the most affected structure. More than 100 genes were upregulated in the amygdala of the 3-month-old AD mice (Figure 5A; Supplementary Table 5), including *Tnfαip8l3, Sstr1, Sstr4, Chrm5* and *Aldh1a3* (Figure 4G; Figure 5A and C). Among the almost 100 downregulated genes in the amygdala at 3 months old (Figure 4G; Supplementary Table 5), we found *Grin3b*, C*hrna10* and *Ciart*. However, when we analysed the number of deregulated genes at 6 months old, both the upregulated and downregulated genes dramatically decreased in all brain areas (Figure 4H, Figure 5B). In amygdala, the upregulation of genes diminished to three genes, *App, Prnp* and *Clec7a* (Figure 4H, Figure 5B, D) while the downregulated genes were restricted to *Dbp* and *Ciart* (Figure 4H; Figure 5C, H).

**Figure 5.**
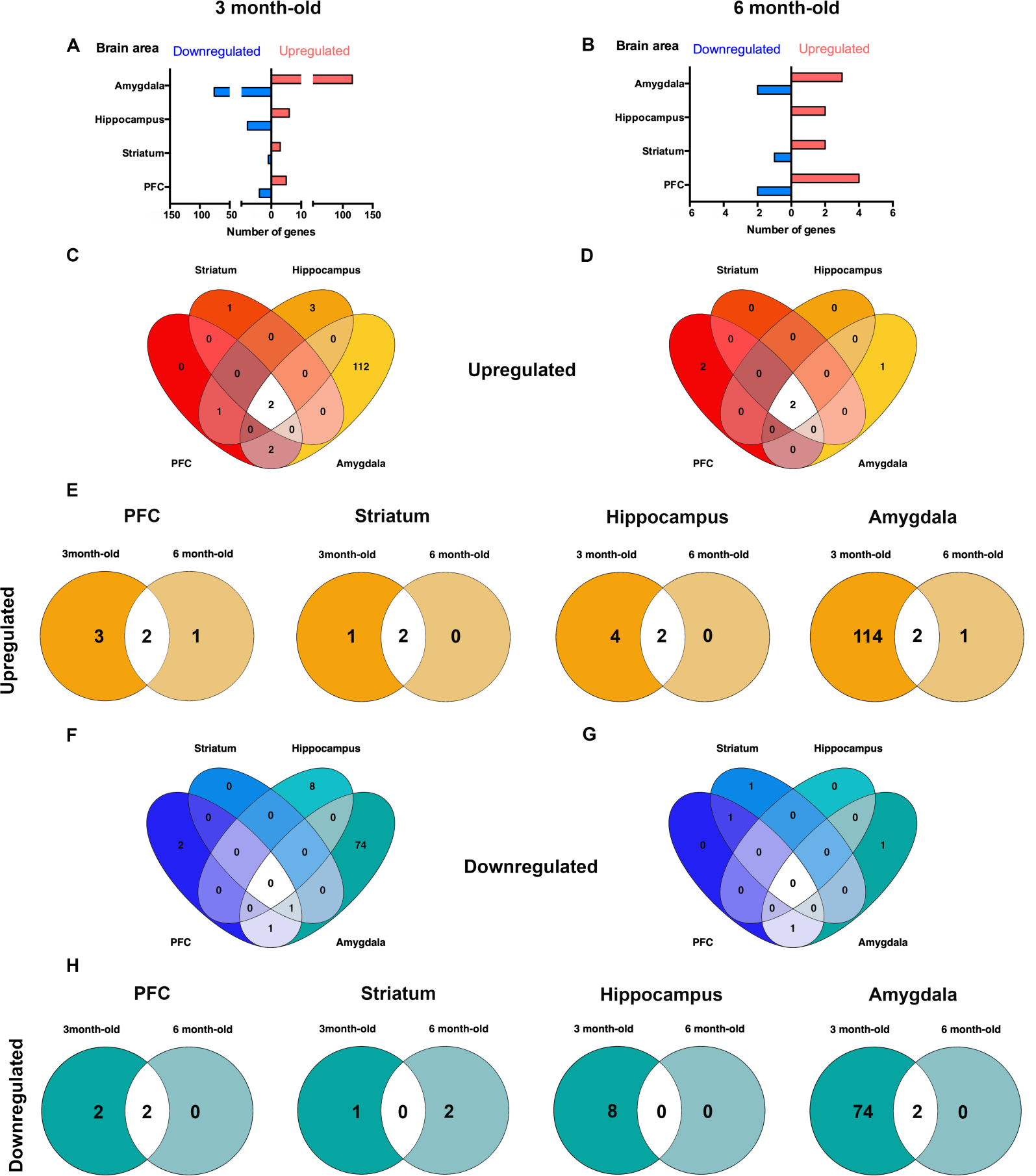
Distribution pattern of differentially expressed genes from APP/PSEN1-Tg mice model. Total number of up (red bars) and downregulated (blue bars) genes within amygdala, hippocampus, striatum and PFC at (A) 3 months and (B) 6 months. Venn diagrams show the number of differentially expressed genes among areas, in an upregulated (C,D) and downregulated (F,G) way. Venn diagrams show the number of upregulated (E) and downregulated (H) shared genes between 3 and 6 months old of age APP/PSEN1-Tg (n=6 per group) *vs* Non-Transgenic mice (n=6 per group).

When we compared those genes that were consistently deregulated at both ages per area, we observed that the upregulation of *App* and *Prnp* were extended throughout PFC, hippocampus and amygdala (Figure 4A,E,G; Figure 5E). Moreover, this upregulation at 6 months-old was affecting all brain regions Figure 4B,D,F and H; Figure 5E). By contrast, *Ciart* was downregulated in PFC, striatum, and amygdala of APP/PSEN1-Tg mice at 3 months old. Only *Nr1d1, Dbp* in PFC and *Dbp* and *Ciart* in amygdala were downregulated in the older transgenic mice (Figure 4B,H; Figure 5H).

### Enriched gene sets at 3 and 6 months-old

We then performed bioinformatic analyses to determine pathways altered in the transgenic mice in the pre-disease and early pathology periods (3 months and 6 months). This functional analysis showed enriched gene sets related to depressive disorders appear at 3 month-old within PFC, hippocampus and amygdala, areas that are intimately involved in both cognitive function and mood regulation (Figure 6A,B,D and E; Supplementary Table 6). In the striatum, we only identified two positively enriched gene sets - those of circadian rhythms and Alzheimer’s disease (Figure 6C) - that also appear in PFC, hippocampus and amygdala at early stages. In the 6 months-old animals, however, the number of gene sets that were significantly enriched in APP/PSEN1-Tg transgenic mice was higher. We identified additional negatively enriched pathways linked to the electron transport chain, found in the PFC, striatum and hippocampus (Figure 6F, G and H; Supplementary Table 7) and also pathways implicated in AD and neuroinflammation. In parallel, the positively enriched depressive-related gene sets not only increased in number but also the dysregulation of these pathways extended to all the brain structures analysed (Figure 6F-I) at this later age. Additionally, the neurotransmission-related pathways were more enriched in 6-month-old transgenic mice than at early ages, suggesting that neurodegeneration increased the number of affected gene sets. By contrast, the impairments in circadian rhythms seem to appear in the preclinical phase in the amygdala (Figure 6B-E), persisting at least to the period when pathology becomes apparent (Figure 6I).

**Figure 6.**
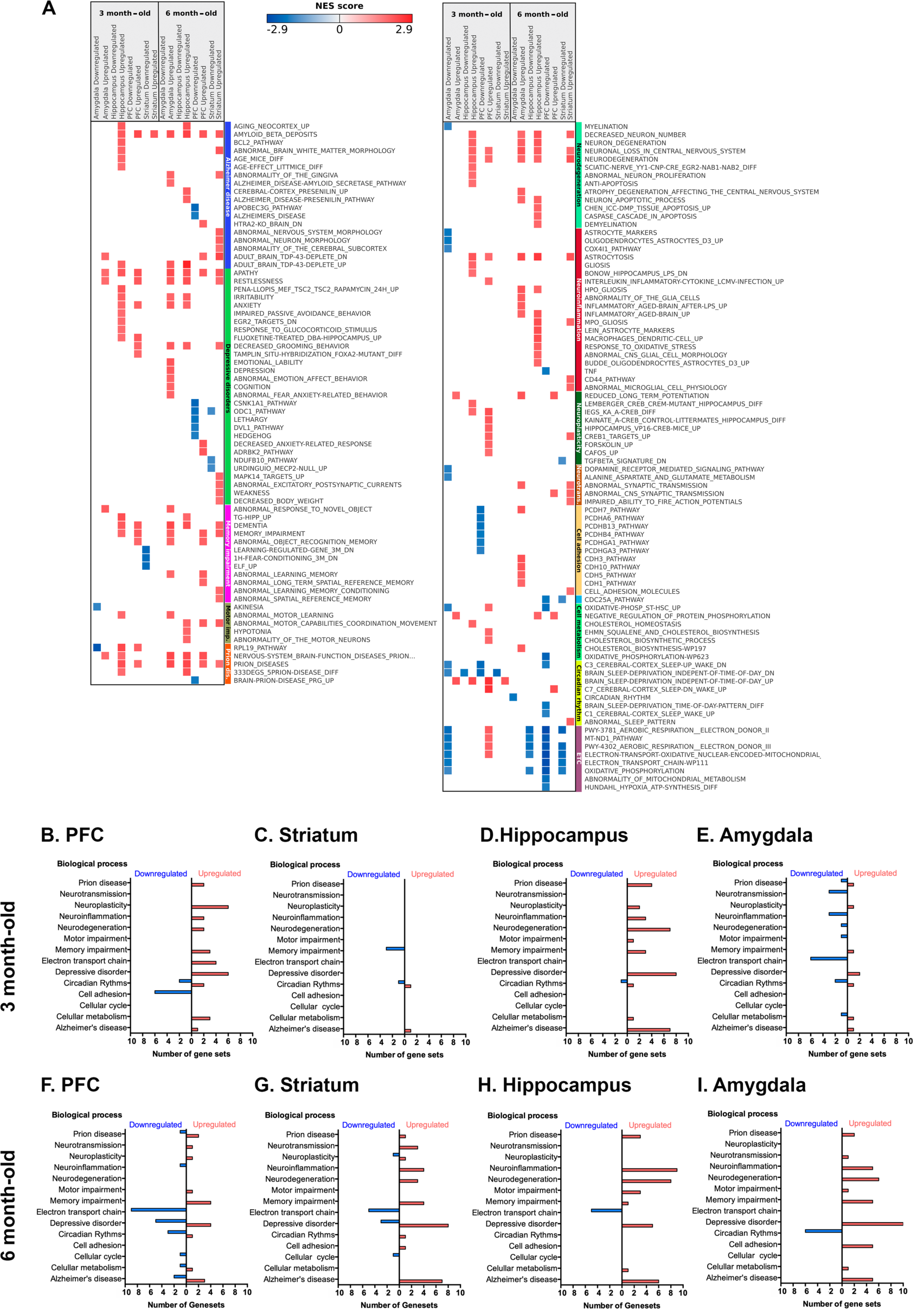
Comparison of differentially expressed gene sets in APP/PSEN1-Tg mice in PFC, striatum, hippocampus and amygdala. (A) The heatmap represents some selected up and downregulated gene sets at 3 and 6 month-old included in different biological processes: Alzheimer disease, depressive disorders, memory impairment, prion disease, motor impairment (Motor imp.), neurodegeneration, neuroinflammation, neuroplasticity, neurotransmission (Neurotrans.), cell adhesion, circadian rhythm, electronic transport chain (ETC). (FDR q value <0.05; |NES| >1.4.). List of the total number of differentially expressed gene sets ordered by biological processes found in PFC (B,F), striatum (C,G), hippocampus (D,G) and amygdala (E,J) at 3 and 6 months old, respectively. Red bars (positive) represent the upregulated expressed gene sets and the blue bars (negative) show downregulated gene sets in each area. For the GSEA we only included those gene sets with FDR q value <0.05 and |NES| >1.4.

We then extended these findings to human data by performing GSEA analysis using DisGeNET (version v7.0; http://www.disgenet.org/) gene sets (Supplementary Figure 3; Supplementary Tables 8 and 9). DisGeNET is a knowledge resource that collects information about the genetic underpinnings of human diseases. This GSEA, which is specific for diseases and disease-related phenotypes, showed positively enriched gene sets corresponding to several subtypes of AD and dementias in the PFC and hippocampus of 3-month-old animals. Additionally, we found positively enriched gene sets associated with memory impairment phenotypes, anxiety disorders, traits and symptoms related to motor impairment, and to neurodegenerative diseases in the hippocampus of younger animals. These results are consistent with the deregulated pathways shown in Figure 6 and with the results of the behavioral tests. In 6-month-old animals, the DisGeNET GSEA showed that positively enriched gene sets of several subtypes of AD and dementias and those related to memory impairment were extended to all brain areas. This dysregulation of processes related to memory impairment in old animals is consistent with the working memory deficits and learning impairments that displayed 6-month-old animals displayed in the behavioural experiments. Moreover, positively enriched gene sets related to diseases of the central nervous system were observed in the hippocampus and striatum of 6-month-old animals. Most of these diseases are cerebrovascular diseases, which have been related to MD^20^. The connection between this class of disorders and MD have been suggested to be mediated by inflammation^21^, which is consistent with the dysregulation of neuroinflammation pathways found in hippocampus of young and old animals (Figure 6A,D,H).

## DISCUSSION

This study provides the first evidence of early, pre-pathological, depression-like symptoms, as well as cognitive impairment, and the underpinning transcriptomic changes specifically in PFC, striatum, hippocampus and amygdala in a rodent AD model (B6C3-Tg (APPswe, PSEN1dD9)85Dbo/MmJax). In particular, APP/PSEN1-Tg transgenic mice show both pre-pathological and peri-pathological deficits in different behaviours. On the one hand, the anxiety-related behaviours and despair behaviours after stressful conditions, hyperlocomotion and diminishing discrimination begins, in the model used, at 3 months of age, before the onset of pathology. On the other hand, the hyposmia traits, working memory deficits and reversal learning impairments appear later at 6 months old, at the time when amyloid plaque pathology begins to become apparent.

The behavioural experiments indicate that APP/PSEN1-Tg mice showed early anxiety-like and depressive-like traits, symptoms which would be expected to be associated with dysfunction of the amygdaloid, hippocampal and PFC circuitries. Supporting these observations, the genes *Nr1d1* and *Bhlhe41*, which have been previously associated with both control of circadian rhythms function and with depression^22,23^, were downregulated in the PFC of transgenic mice. Moreover, *Plch1* is upregulated at 3 months old in the transgenic mice. The product of this gene is part of the pathway generating second messengers, such as inositol 1,4,5-triphosphate and diacylglycerol; a pathway which is intimately linked to the depressive-like phenotype^24^. At 3 but not at 6 months old, we found the upregulation of *Inhba* in PFC and hippocampus, which could represent a compensatory change to mitigate the depressive-like behaviour. In fact, the activin signalling pathway plays a key role in the response to antidepressant treatment in humans^25^. *Nr1d1* remains downregulated at 6 months old, without changes in *Inhba* signalling, due to possible deeper detrimental effects at older ages, e.g. deficits in working memory. Additionally, *Mc4r* is downregulated in hippocampus at 3 months old and facilitates anxiety-like and depression-like behaviours pursuant to chronic stress^26^. Indeed, a general overexpression of *Prnp* and *App* in 3- and 6-month-old APP/PSEN1-Tg mutant mice could also contribute to this depressive-like symptomatology, since previous studies found alterations of these protein levels in cortical areas and peripheral blood of MD^27,28^. Noteworthy, this mouse overexpresses APP695 containing the double Swedish mutation, so the general upregulation of *App* found in all brain areas of interest is directly induced by the model. Moreover, *Prnp* may modulate depressive-like behaviour in mice via interactions with monoaminergic neurotransmission^29^. In accordance with these results, we observed an increasing number of enriched pathways linked to depressive disorders (including depression, apathy and anxiety, among others) and disease-associated gene sets at 3 and 6 months old, before the period of formation of β-amyloid plaques in this model^30^. These results therefore support the notion that depressive symptoms are part of the prodromal phase of the AD in this rodent model, just as they are hypothesised to be in human disease.

Overall, the APP/PSEN1-Tg mutant mice had a general decline in memory with ageing, as previously described in AD models^15,31^ but these phenomenon was demonstrated before the onset of pathology, although became more extensive at about the time pathology was expected. Despite a subtle imbalance of memory impairment pathways at 3 months old, transgenic mice showed higher latencies to acquire the criterion in the left-right discrimination test. These memory deficits could be promoted by the observed dysregulation of *Cdh23, Ptx4* and *Mc4r* genes in hippocampus, as they are known to participate in the endocytosis of glutamatergic ionotropic AMPA receptors, hippocampal synaptic plasticity and mood-like disorders in mice, respectively^32–35^. At 6 months old, APP/PSEN1-Tg mutant mice show deeper deficits in memory in all the evaluated tasks. They display longer latencies in left-right discrimination task, and deficits in working and long-term memories. These memory impairments run in parallel with an increasing number enriched pathways in PFC and hippocampus (abnormal long-term spatial reference and spatial memories, among others), involved in working^36,37^ and spatial learning^38,39^, respectively, and with dysregulation of gene sets related to memory impairment and learning disabilities (Supplementary Figure 3).

Our results show that the APP/PSEN1-Tg, at 3 months old mice spent the same time than the control mice investigating the cotton-swabs, which have been impregnated with limonene. However, at 6 months old the exploration of both odours was increased in transgenic mice compared to the control group, independently whether water and limonene were presented. Overall, APP/PSEN1-Tg animals seem not to present a complete anosmia, although it seems that they display a loss of discrimination between odours, so this possible hyposmia together with working memory^36,37^ impairments could explain the indiscriminate investigation that transgenic mice display. These olfactory alterations could be due to pathology in the olfactory bulb at 6 months-old^40^, that might be promoting changes in the sense of smell of APP/PSEN1-Tg mice. Thus, our mice model could reflect the progressive anosmia that AD patients show at early stages of the disease^19^. Indeed, a recent study shows that late-depression patients show similar olfactory impairments that those in AD^41^. In this sense, early olfactory deficits could lead to the identification of depression patients with high risk of developing AD^41^, as also occurs in our APP/PSEN1-Tg animals by showing early depression-like behaviour with later hyposmia traits.

MD and AD are both associated with neuroinflammation and oxidative stress^11^ and evidence for the activation of both processes was observed in the differential expression analysis. This show a dysregulation of inflammatory-related response genes, such as *Maml3, Zap70, Itgax* and *App*^42–45^ at both ages in APP/PSEN1-Tg mice. In addition, gene sets related to electron transport chain are, in general, downregulated in APP/PSEN1-Tg animals at 6 months in PFC, striatum and hippocampus, reflecting a possible abnormal function of mitochondrial metabolism, as previously observed in AD^46^ and MD^47^.

One of the most striking observations is that majority of the negatively enriched gene sets are associated with control of circadian rhythms, and these are observed in all brain areas of interest (Figure 6B). In particular, we observe the downregulation of *Ciart* and *Dbp* in amygdala, striatum and PFC at 3 and 6 months old (Figure 5 for more details). The expression of both genes in anterior cingulate cortex modulates the circadian rhythms and mood^23,48^, through the MERK/ERK signalling pathway. Although disturbances in the circadian rhythms are associated with both MD and AD^49^, this is the first study in which both circadian transcripts are linked to AD. A close relationship has been found between *Dbp* and AD, given that TGF-β2, present in the cerebrospinal fluid of AD patients, inhibits the expression of some clock genes, including *Dbp*, in *in vitro* assays. We also note that the predominant tau-kinase, reported to be downstream of β-amyloid production^50–52^, GSK3β, is a regulator of Per2 phosphorylation and hence a master-regulator of the mammalian clock^53^. Our results suggest that there is a decline in the functional control of circadian rhythms in APP/PSEN1-Tg mice that should be further studied in detail.

In summary, we suggest that MD could be an indicator of the early onset of AD. The APP/PSEN1-Tg model of AD could recreate the early-depressive symptoms before developing AD, as a prodromal stage of the neurodegenerative disease. In fact, this model provides *App, Prnp, Ciart* and *Dbp* as possible shared biomarkers of AD and MD, despite the implications of *App* and *Prnp* in MD remaining unclear^54–56^. In this sense, further studies combining post-mortem human tissues of PFC, striatum, hippocampus and amygdala from AD patients with and without an early history of depression, and transcriptomic analysis could help to provide additional evidence on the combined biological signature of both diseases.

## MATERIAL AND METHODS

See Supplementary material and methods for additional details.

### Animals and rearing conditions

For the present study, we used 30 hemizygous double transgenic male mice (B6C3-Tg (APPswe, PSEN1dD9)85Dbo/MmJax) model of AD (APP/PSEN1-Tg) and 30 male Non-Transgenic control mice (004462, The Jackson Laboratory, USA). These transgenic mice express a chimeric mouse/human APP (Mo/HuAPP695swe) and a mutant human presenilin-1 (PS1-dD9), each controlled by independent mouse prion protein promoter elements^30^. The mice were assigned to two different groups: APP/PSEN1-Tg and Non-Transgenic (n=16 per group) at 3 months-old, and APP/PSEN1-Tg and Non-Transgenic (n=14 per group) at 6 month-old, before developing the β-amyloid plaques and right at onset^30^. All procedures were conducted in accordance with national (BOE-2013-1337) and EU (Directive 2010-63EU) guidelines regulating animal research and were approved by the local ethics committee (CEEA-PRBB).

### Behavioural Evaluation

*Elevated plus maze* (EPM) (Panlab s.l.u, Barcelona, Spain) was performed using a black maze elevated 30 cm above the ground^57^. Each mouse was placed in the centre of the maze, and was allowed to freely explore for 5 min. The software SMART (Panlab s.l.u., Spain) automatically recorded the number of entries and the time spent in the arms.

### Novel object recognition (NOR)

Single trial NOR was performed in an open black arena (32 × 28 cm) using 3 object types at opposite corners of the open field, 50 mm from the walls, similar to those previously described^58^. Object exploration was defined as intentional contact between the mouse’s nose and the objects. The recognition index was defined as [t_Novel_/(t_Novel_ + t_Familial_)] x 100 for animals exploring novel objects in the acquisition trial.

### Left-right discrimination learning

This test was performed using a T-maze apparatus, as previously described^31^. This T-maze was filled with water (23 ± 1 °C). During the first two trials, two identical platforms were submerged on the end of both arms to test possible side preferences. A mouse was considered to have achieved criterion after 5 consecutive errorless trials. The reversal-learning phase was then conducted 48h later, applying the same protocol except that mice were trained to reach the escape platform of the opposite arm. Escape latencies and number of trials to reach the criterion were manually recorded. A maximum of 20 trials were needed to complete the experiment.

#### Tail suspension test (TST)

Each mouse was suspended individually 50 cm above a bench top for 6 min^57^. Mice were individually video-recorded and an observer, blind to the experimental conditions, evaluated the percentage of time the animal was immobile during the test.

#### Exposure to stress

The stressful procedure consisted in the exposition to two mild stressful situations each day during four consecutive days: animals were placed for 10 min in the open field apparatus, and then they were placed in glass cylinders filled with water during 6 min. Animals were subsequently evaluated for TST.

#### Habituation-dishabituation test

The test was performed as described^59^. After five minutes of habituation to the cage, six consecutive one-minute presentations of a cotton swab with distilled water were followed by one presentation of limonene (Sigma-Aldrich) solution 1:1000 in distilled water. After that, five consecutive one-minute applications of distilled water were presented. Tests were video-recorded and a blinded observer measured the time that mice spent sniffing the cotton tip rearing on its hind limbs.

#### Animal sacrifice and sample preparation for RNA extraction

After behavioural analysis, animals were sacrificed by cervical dislocation and brains were immediately removed from the skull. Brain samples were dissected at both ages: 3 and 6 months old from Non-Transgenic and APP/PSEN1-Tg mice (n=6 per group). PFC (Figure 4I), striatum, amygdala and hippocampus were dissected following an anatomical atlas^60^. Brain samples were immediately stored at -80°C until the RNA extraction. Then, each tissue sample was homogenized in 1ml of QIAzol Lysis Reagent using a rotor–stator homogenizer (Polytron PT 2500 E; Kinematica AG, Switzerland) during 20–40 seconds. After homogenization, the RNA was extracted using RNeasy Lipid Tissue Mini Kit (Qiagen)^61^.

### RNA sequencing

#### Library preparation and sequencing

RNA samples (50-100 ng) with RIN scores from 7.6 to 9 (Agilent 4200 TapeStation) were reverse transcribed to cDNA. Poly-A tail selection was done using Total Dual RNA-Seq PolyA. cDNA was sequenced (HiSeq3000/4000) at the Oxford Wellcome Trust facility obtaining 75 bp per read. On average, 33M reads were obtained per sample and a mapping rate of ∼85%.

### RNAseq DE analysis

Raw sequencing reads in the 1710 paired fastq files were mapped with STAR version 2.5.3a^62^ to the Gencode release 17 based on the GRCm38.p6 reference genome and using the corresponding GTF file. The 885 bam files corresponding to 9 different lanes were merged for each sample using Samtools 1.5, ending up with 95 bam files. The table of read counts was obtained with featureCounts tool in subread package, version 1.5.1.

Further analyses were performed in R, version 3.4.3. Genes having less than 10 counts in at least 10 samples were excluded from the analysis. Raw library size differences between samples were treated with the weighted “trimmed mean method” TMM^63^ implemented in the edgeR package^64^. These normalized counts were used in order to make the unsupervised analysis, heatmaps and clusters. For the differential expression (DE) analysis, read counts were converted to log2-counts-per-million (logCPM), the mean-variance relationship was modelled with precision weights using the voom approach and linear models were subsequently applied with limma package, version 3.30.13^65^. p-values (p) were adjusted for multiple comparisons using the Benjamini-Hochberg false discovery rate (FDR) approach. Genes were considered to be differentially expressed if |logFC|>0.585 and adjusted p<0.05.

### qPCR validation

Data were obtained from the qPCR platform using Taqman Low Density array (TLDA; qPCR 7900HT, Life Technologies), where thirteen genes of interest were studied together with the following endogenous controls: *Gapdh, Tbp* and *18S*. The gene *18S* was used to check overall expression whereas *Gapdh* and *Tbp* were geometric averaged and included in posterior analyses as the *C*_*t*_*(endogenous)*. For each gene, DC_t_ was computed, comparing these values between conditions. *DC*_*t*_*= C*_*t*_*(gene) - C*_*t*_*(endogenous)*. Comparisons between studied conditions were performed using a t-Student’s t-test. Results were adjusted for multiple comparisons using the FDR. The comparisons performed in each area are APP/PSEN1-Tg and Non-Transgenic at both ages (validated genes in Supplementary Table 1).

### Functional and Pathway analysis

Pre-ranked GSEA was used in order to retrieve enriched gene sets corresponding to functional pathways^66^. The list of genes was ranked using the -log(p.val)*signFC value for each gene from the statistics obtained in the DE analysis with limma.

For the gene set collection we used a database described previously^67^ and available in http://ge-lab.org/gs/. The gene sets in this database are harvested from different pathway data sources and published studies all related to mice.

MGI genes IDs were converted to mouse symbol ID using the annotation files in (http://www.informatics.jax.org/). As there were a high number of redundant gene sets in the collection, a filtering strategy was applied to select the most representative ones. The filtering steps were: i) pairwise Jaccard coefficients were computed between the gene sets from the following data sources: Published articles, MPO, HPO, Reactome, MsigDB, GO_BP, KEGG, PID, WikiPathways, INOH, PANTHER, NetPath, Biocarta, EHMN, MouseCyc and HumanCyc; ii) for gene sets with a Jaccard = 1 (gene sets constituted by the same genes), only one was considered, and iii) from gene sets with a Jaccard > 0.8, the gene set with the highest number of genes was considered and the other gene sets were excluded from the gene set collection.

### DisGeNET database analysis

We performed a GSEA to retrieve enriched gene sets associated with human diseases. For this, we ranked the genes using the -log(p.val)*signFC value from the DE analysis. The gene sets associated with human diseases, symptoms, and traits were retrieved from the DisGeNET database (https://www.disgenet.org/, v7)^68^. DisGeNET integrates gene-disease associations collected from curated repositories, from genome-wide association studies (GWAS) catalogues, from animal models, and data obtained from the scientific literature using text mining approaches^68^. For this analysis, we used only the curated gene sets in DisGeNET, that include information from repositories such as UniProt, PsyGeNET, Clingen, and the Genomics England Panel app. The mouse genes were converted to human gene IDs using the annotation files in (http://www.informatics.jax.org/).

## STATISTICAL ANALYSIS

For animal model experiments, analysis was performed using software SPSS 23.0 (SPSS Inc., USA). Statistical differences were investigated by Student’s t-test and ANOVA with repeated measurement analysis. *Post-hoc* analysis was calculated with Bonferroni’s correction when applicable. Since habituation-dishabituation data were not normally distributed, Wilcoxon matched-pairs rank test and Kruskal-Wallis test were calculated. Fisher’s exact test was used to evaluate differences in the percentage of animals that made an error in the Y-maze. For the GSEA we only included those gene sets with a FDR q value<0.05 and |NES| >1.4.

## DATA AVAILABILITY

The data that support the findings of this study are available from the corresponding author upon reasonable request.

## ABBREVIATIONS

AD: Alzheimer’s disease
APP: Beta-Amyloid precursor protein
BSPD: Behavioural and psychological symptoms of dementia
DE: Differential expression
EPM: Elevated plus maze
FC: Fold change
FDR: False discovery rate
GSEA: Gene set enrichment analysis
GWAS: Genome-wide association study
ICD-9: International Classification of Diseases, 9^th^ Edition
MD: Major Depression
NES: Normalized enrichment score
NOR: Novel object recognition
OF: Open field
PFC: Prefrontal cortex
PSEN1: Presenilin-1
TST: Tail suspension test

## FUNDING AND DISCLOSURE

This study was funded by the UE Medbioinformatic project (grant number 634143), Ministerio de Economia y Competitividad (grant number SAF2016-75966-R-FEDER), Ministerio de Sanidad (Retic-ISCIII, RD16/017/010 and Plan Nacional sobre Drogas 2018/007). The Department of Experimental and Health Sciences (UPF) is a “Unidad de Excelencia María de Maeztu” funded by the AEI (CEX2018-000792-M). The Research Programme on Biomedical Informatics (GRIB) is a member of the Spanish National Bioinformatics Institute (INB), funded by ISCIII and FEDER (PT17/0009/0014). The GRIB is also supported by the Agència de Gestió d’Ajuts Universitaris i de Recerca (AGAUR), Generalitat de Catalunya (2017 SGR 00519).

## AKNOWLEDGEMENTS

The authors are indebted to the Oxford Genomics Centre at the Wellcome Centre for Human Genetics (funded by Wellcome Trust grant reference 203141/Z/16/Z) for the generation and initial processing of sequencing data. The authors wish to thank Adriana Castro-Zavala for her expertise and collaboration in tissue extraction for the RNA sample preparation.

## AUTHOR CONTRIBUTIONS

A.M-S, L.N, L.I.F., F.S. and O.V. were responsible for the study concept and design. A.M-S. performed behavioural studies. A.M-S, E.M.R., A.H.N. and S.L. contributed to transcriptomic analysis. J.P., M.A., L.N., F.S. and L.I. F. carried out the bioinformatics analysis. All the authors participated in the interpretation of findings. A.M-S., F.S., and O.V. drafted the manuscript. All authors critically reviewed the content and approved the final version for publication.

## CONFLICTS OF INTEREST

The authors declare no conflicts of interest regarding the work presented here.

## Notes

### Competing Interest Statement

The authors have declared no competing interest.

